# Consequences of natural perturbations in the human plasma proteome

**DOI:** 10.1101/134551

**Authors:** Benjamin B. Sun, Joseph C. Maranville, James E. Peters, David Stacey, James R. Staley, James Blackshaw, Stephen Burgess, Tao Jiang, Ellie Paige, Praveen Surendran, Clare Oliver-Williams, Mihir A. Kamat, Bram P. Prins, Sheri K. Wilcox, Erik S. Zimmerman, An Chi, Narinder Bansal, Sarah L. Spain, Angela M. Wood, Nicholas W. Morrell, John R. Bradley, Nebojsa Janjic, David J. Roberts, Willem H. Ouwehand, John A. Todd, Nicole Soranzo, Karsten Suhre, Dirk S. Paul, Caroline S. Fox, Robert M. Plenge, John Danesh, Heiko Runz, Adam S. Butterworth

## Abstract

Proteins are the primary functional units of biology and the direct targets of most drugs, yet there is limited knowledge of the genetic factors determining inter-individual variation in protein levels. Here we reveal the genetic architecture of the human plasma proteome, testing 10.6 million DNA variants against levels of 2,994 proteins in 3,301 individuals. We identify 1,927 genetic associations with 1,478 proteins, a 4-fold increase on existing knowledge, including *trans* associations for 1,104 proteins. To understand consequences of perturbations in plasma protein levels, we introduce an approach that links naturally occurring genetic variation with biological, disease, and drug databases. We provide insights into pathogenesis by uncovering the molecular effects of disease-associated variants. We identify causal roles for protein biomarkers in disease through Mendelian randomization analysis. Our results reveal new drug targets, opportunities for matching existing drugs with new disease indications, and potential safety concerns for drugs under development.

## Introduction

Plasma proteins play key roles in biological processes such as signalling, transport, growth, repair, and defence against infection. They are frequently dysregulated in disease and are the targets of many drugs. Detailed characterisation of the genetic factors that determine inter-individual protein variability will, therefore, furnish both fundamental and applied insights^1^. Despite evidence of the heritability of plasma protein abundance^2^, systematic genome-wide study of such ‘protein quantitative trait loci’ (pQTLs) has been constrained by inability to measure large numbers of proteins reliably in large numbers of individuals^1,3–5^.

Here we combine several technical and conceptual advances to create and interrogate a rich genetic atlas of the human plasma proteome. First, we use a markedly expanded version of an aptamer-based multiplex protein assay (SOMAscan)^6^ to quantify 3,620 plasma proteins in 3,301 healthy individuals, representing several-fold increases both in breadth of protein panel and in cohort size. Second, we exploit improvements in genotype imputation panels to achieve 10-fold denser genotypic coverage than in previous proteomic studies^7^. Third, we draw on studies with multi-dimensional data (e.g., transcriptomics and clinical phenotypes) and on new bioinformatics tools to help understand mechanisms and clinical consequences of perturbations in protein pathways^8^. Fourth, we use two-sample “Mendelian randomization” techniques to evaluate the causal relevance of protein biomarkers to disease^9^. Finally, we cross-reference genomic-proteomic information with disease and drug databases to identify and prioritise therapeutic targets.

Our study characterises the genetic architecture of the human plasma proteome, identifying 1,927 genotype-protein associations, including *trans-*associated loci for 1,104 proteins that provide key insights into protein regulation. More than 150 pQTLs overlap with disease susceptibility loci, elucidating the molecular effects of disease-associated variants. We find strong evidence to support causal roles in disease for several protein pathways, highlighting novel therapeutic targets as well as potential safety concerns for drugs in development.

### Genetic architecture of the plasma proteome

After stringent quality control, we performed genome-wide testing of 10.6 million autosomal variants against levels of 2,994 plasma proteins in 3,301 healthy European-ancestry individuals (Methods, Extended Data Figure 1). Genotypes were measured using the Affymetrix Axiom UK Biobank array and imputed against a combined 1000 Genomes and UK10K reference panel. Protein levels were measured using the SOMAscan assay. We evaluated the robustness of protein measurements in several ways (Methods, Supplementary Note). Measurements in replicate samples were highly consistent; the median coefficient of variation across all proteins was 0.064 (interquartile range 0.049-0.092). We also showed temporal consistency in protein levels in samples obtained two years apart from the same individuals, reproduced known associations with non-genetic factors, and verified selected protein measurements with multiple different assay methods.

We found 1,927 genome-wide significant (*p*<1.5×10^−11^) associations between 1,478 proteins and 764 genomic regions (Figure 1a, Supplementary Table 1, Supplementary Video 1), with 89% of pQTLs reported here for the first time. Of these 764 regions, 502 (66%) had *cis* associations only, 228 (30%) had *trans* associations only and 34 (4%) had both *cis* and *trans*. 95% and 87% of our *cis* pQTL variants were located within 200Kb and 100Kb, respectively, of the relevant gene’s transcription start site (TSS) (Figure 1b), and 44% were within the gene itself. The *p*-values for *cis* pQTL associations increased with increasing distance from the TSS, mirroring findings for *cis* expression QTLs (eQTLs) in transcriptomic studies^10,11^. Of the proteins for which we identified a pQTL, 88% had either *cis* (n=374) or *trans* (n=925) associations only, while 12% (n=179) had both. The majority of significantly associated proteins (75%; n=1,113) had a single pQTL, while 20% had two and 5% had more than two (Figure 1c). To detect multiple signals at the same locus we used stepwise conditional analysis, identifying 2,658 conditionally significant associations (Supplementary Table 2). Of the 1,927 locus-protein associations, 414 (21%) had multiple conditionally significant signals (Figure 1d), of which 255 were *cis*.

**Figure 1.**
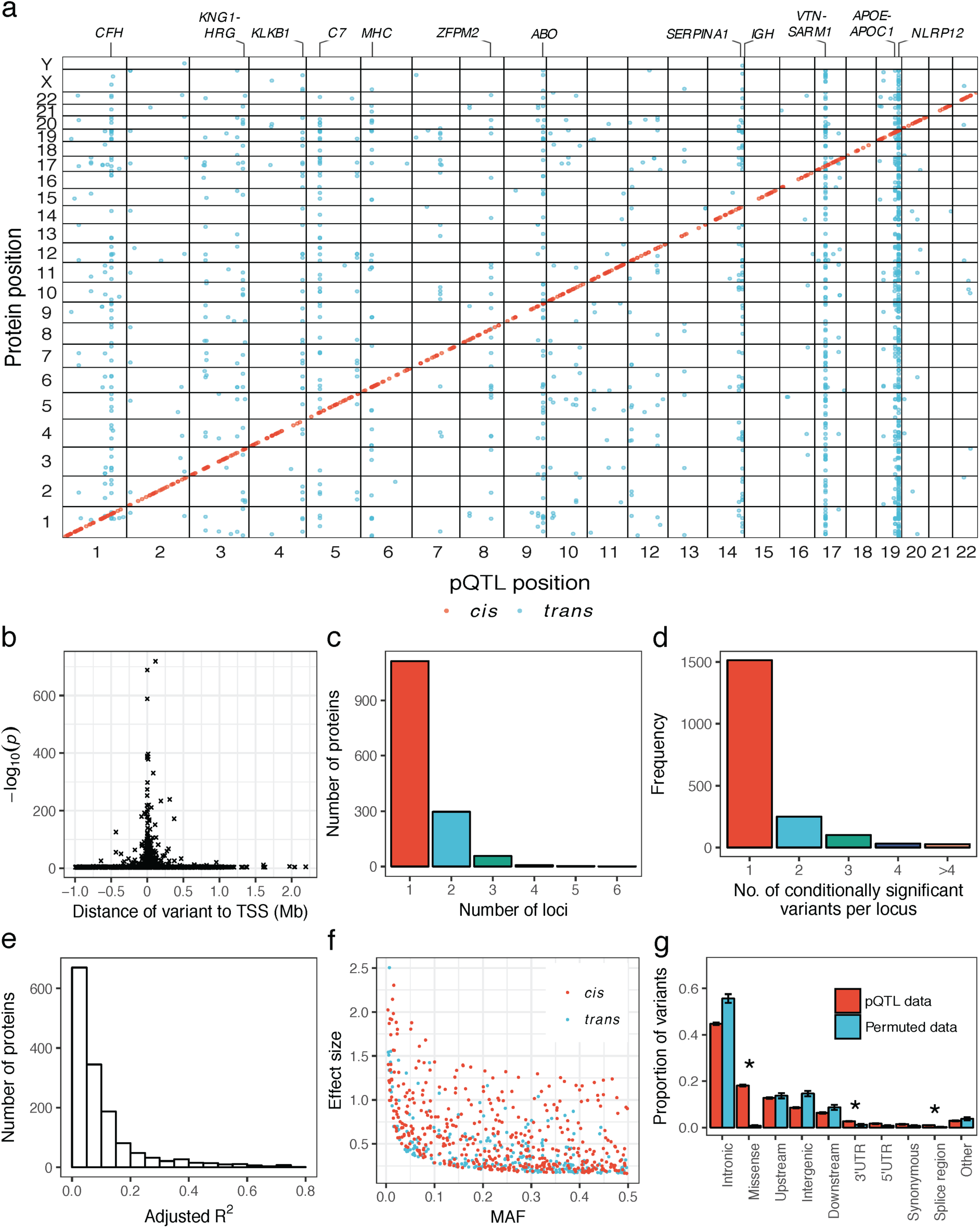
The genetic architecture of plasma protein levels. (a) Genomic location of pQTLs. Plot of sentinel variants for pQTLs (red= *cis*, blue= *trans*). Y-axis indicates the position of the gene that encodes the associated protein. The twelve most associated regions of the genome are annotated. (b) Plot of the statistical significance of the most associated (sentinel) *cis* variant for each protein against the distance from the transcription start site (TSS). (c) Histogram of the number of significantly associated loci per protein. (d) Histogram of the number of conditionally significant signals within each associated locus. (e) Histogram of protein variance explained (adjusted R^2^) by conditionally significant variants. (f) Distribution of effect size against minor allele frequency (MAF) for *cis* and *trans* pQTLs. (g) Distribution of the predicted consequences of the sentinel pQTL variants compared to matched permuted null sets of variants. Asterisks highlight empirical enrichment *p*<0.005.

Genetic variants that change the protein structure may result in apparent pQTLs due to altered aptamer binding rather than true quantitative differences in protein levels. Using bioinformatic approaches to evaluate the extent to which *cis* pQTLs might reflect technical effects (Methods) we found that 91% of pQTLs were unlikely to be influenced by differential aptamer binding (Supplementary Tables 1 and 3). Furthermore, in sub-studies that involved immunoassays, we found strongly concordant genotype-protein associations (Supplementary Note). Even where apparent pQTLs do arise from alternative protein structure that may affect aptamer binding rather than differences in protein abundance, such signals may be functionally significant.

The median variation in protein levels explained by our pQTLs was 5.8% (in-sample estimate; interquartile range: 2.6%-12.4%, Figure 1e). For 193 proteins, however, genetic variants explained more than 20% of the variation, such as for teratocarcinoma-derived growth factor 1 (65%) and haptoglobin (55%). We found a strong inverse relationship between effect size and minor allele frequency (MAF) (Figure 1f), consistent with previous genome-wide association studies (GWAS) of quantitative traits^12–14^. We found 25 associations with rare (MAF <1%) variants and a further 207 associations with low-frequency (MAF 1-5%) variants. Of the 31 strongest pQTLs (per-allele effect size >1.5 standard deviations), 25 were rare or low-frequency variants.

pQTLs were strongly enriched for missense variants (*p*<0.0001) and for location in 3’ untranslated (*p=*0.0025) or splice regions (*p=*0.0004) (Figure 1g, Extended Data Figure 2). To assess whether pQTLs were enriched within regulatory elements from a wide range of cell types and tissues^15–17^, we used GARFIELD^18^ (Methods). We found strong (≥ 3-fold, *p*<5×10^−5^) enrichment of pQTLs in blood cells – unsurprisingly given our use of plasma - at features indicative of transcriptional activation (Extended Data Figure 3, Supplementary Table 4). We also found enrichment at hepatocyte regulatory elements, consistent with the liver’s role in producing many secreted proteins.

### Overlap of loci for gene expression and protein levels

A fundamental biological question is the extent to which genetic associations with plasma protein levels are driven by effects at the transcription level, rather than other mechanisms, such as altered protein clearance or cleavage of membrane receptor proteins from the cell surface. We therefore cross-referenced our *cis* pQTLs with a wide range of publicly available eQTL studies involving >30 tissues or cell types (Supplementary Table 5) using PhenoScanner^8^. 40% (n=224) of *cis* pQTLs had an association with expression of the same gene in at least one tissue (Supplementary Table 6), consistent with similar comparisons within lymphoblastoid cell lines (LCLs)^19^. The greatest overlaps were found in whole blood (n=117), liver (n=70) and LCLs (n=52), consistent with biological expectation, but also likely driven by the larger sample sizes for expression studies of these cell types. *Cis* pQTLs were significantly enriched (*p<*0.0001) for eQTLs for the corresponding gene (mean 20% versus 2% in a background permuted set; Methods, Supplementary Table 7). To address the converse (i.e., to what extent do eQTLs translate into pQTLs), we used a subset of well-powered eQTL studies in relevant tissues (whole blood, LCLs, liver and monocytes^20–23^). Of the strongest *cis* eQTLs (*p<*1.5×10^−11^), 12.2% of those in whole blood were also *cis* pQTLs, 21.3% for LCLs, 14.8% for liver and 14.7% for monocytes.

Comparisons between eQTL and pQTL studies have inherent limitations due to differences in the tissues, sample sizes and technological platforms used. Moreover, plasma protein levels may not reflect levels within tissues or cells. Nevertheless, our data suggest that plasma protein abundance is often, but not exclusively, driven by regulation of mRNA. Our finding that pQTLs are enriched in gene regulatory regions supports the notion that transcription has a major role. *Cis* pQTLs without corresponding *cis* eQTLs may be underpinned by effects not captured at the mRNA level, such as differences in protein stability, degradation, binding, secretion, or clearance from circulation.

### Using *trans* pQTLs to illuminate biological pathways

*Trans* pQTLs are particularly useful for understanding biological relationships between proteins if a causal gene at the *trans*-associated genetic locus can be identified. To this end, we used a combination of databases of molecular pathways and protein-protein interaction networks, and functional genomic data that include variant annotation, eQTL and chromosome conformation capture, to link *trans*-associated variants to potential causal genes (Methods, Supplementary Table 8, Extended Data Figure 4). Of the 764 protein-associated regions, 262 had *trans* associations with 1,104 proteins. We replicated previously reported *trans* associations including *TMPRSS6* with transferrin receptor protein 1^24^ and *SORT1* with granulins^25^. Most (82%) *trans* loci were associated with fewer than four proteins. However, we observed twelve regions with more than 20 associated proteins (Figure 1a, Extended Data Figure 5), including well-known pleiotropic loci (e.g., *ABO*, *CFH*, *APOE*, *KLKB1*) and loci associated with many correlated proteins (e.g., the locus containing the transcription factor gene, *ZFPM2*).

We identified a number of novel associations with strong biological plausibility (Supplementary Table 9, Supplementary Note). Growth differentiation factor 8 (GDF8, more commonly known as myostatin) provides one such example. Because insufficient myostatin has been shown to result in excessive muscle growth^26^, myostatin inhibition has emerged as a promising therapeutic strategy for treatment of conditions characterised by muscle weakness, such as muscular dystrophy^27^. We identified a common allele (rs11079936:C) near *WFIKKN2* that is associated in *trans* with lower levels of plasma GDF11/8 (*p*=7.9×10^−12^), as well as in *cis* with lower levels of plasma WFIKKN2 (*p*=6.9×10^−136^, Supplementary Table 1, Figure 2). The *trans* association attenuated completely upon adjustment for levels of WFIKKN2 (*p=*0.7), while the *cis* association remained significant after adjustment for GDF11/8 (*p*=7.2×10^−113^), suggesting that WFIKKN2 regulates GDF11/8. This observation is supported by *in vitro* evidence suggesting that WFIKKN2 has high affinity for GDF8 and GDF11^28^. This finding illustrates how delineation of causal genes at *trans-*associated loci may help identify additional drug targets in pathways known to be relevant to disease.

**Figure 2.**
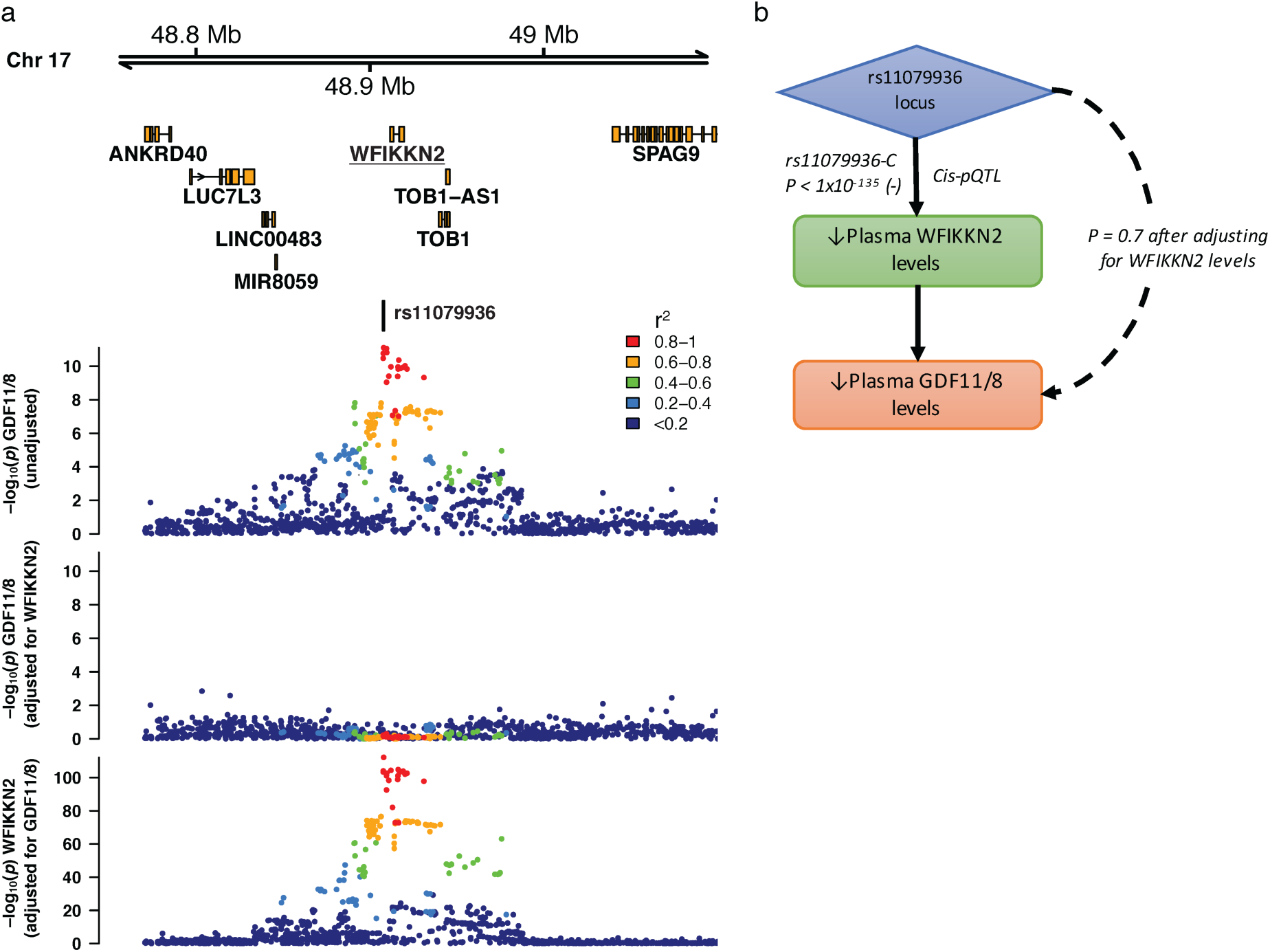
The GDF11/8 *trans* pQTL is mediated by genetic control of WFIKKN2 levels. (a) Regional association plots of the *trans* pQTL (sentinel variant rs11079936) for GDF11/8 before and after adjusting for levels of WFIKKN2 (upper panels), and the WFIKKN2 *cis* pQTL after adjusting for GDF11/8 levels (bottom panel). A similar pattern of association for WFIKKN2 was seen prior to GDF11/8 adjustment (not shown). (b) Proposed mechanism of how the *trans* pQTL for GDF11/8 is mediated by WFIKKN2 levels.

### Identifying causal pathways underlying disease susceptibility loci

GWAS have identified thousands of loci associated with common diseases, but the pathways by which most variants influence disease susceptibility await discovery. To identify intermediate links between genotype and disease, we overlapped pQTLs with disease-associated genetic variants identified through GWAS (*p*<5×10^−8^). 152 of our pQTLs were strongly correlated (*r*^2^ ≥ 0.8) with variants significantly associated with disease (Supplementary Table 10), including 38 with *cis* associations, 109 with *trans* associations and 5 with both. In the examples below, we illustrate how our findings provide novel insights spanning a wide range of disease domains including autoimmunity, cancer and cardiovascular disease.

Our *trans* pQTL data implicate previously unsuspected proteins as mediators through which genetic loci exert their influence on disease risk. For example, GWAS have identified a missense allele (rs3197999:A, p.Arg703Cys) in *MST1* on chromosome 3 that increases risk of inflammatory bowel disease (IBD) (Figure 3)^29,30^, and decreases plasma MST1 levels^31^. We show that this polymorphism acts in *trans* to reduce abundance of BLIMP1, encoded by *PRDM1* on chromosome 6 (Figure 3, Supplementary Table 1). BLIMP1 is a transcriptional master regulator that plays a critical role in the terminal differentiation of immune cells. Intriguingly, there is another IBD association signal in the intergenic region adjacent to *PRDM1* on chromosome 6 (Figure 3)^30^. Both *PRDM1* and its neighbour *ATG5*, which encodes a protein involved in autophagy, are plausible candidate genes at this locus. Our data implicate BLIMP1 as a previously unidentified mediator of the IBD association in *MST1* on chromosome 3. In addition, our results provide indirect support for the hypothesis that *PRDM1* is the causal gene underlying the chromosome 6 association.

**Figure 3.**
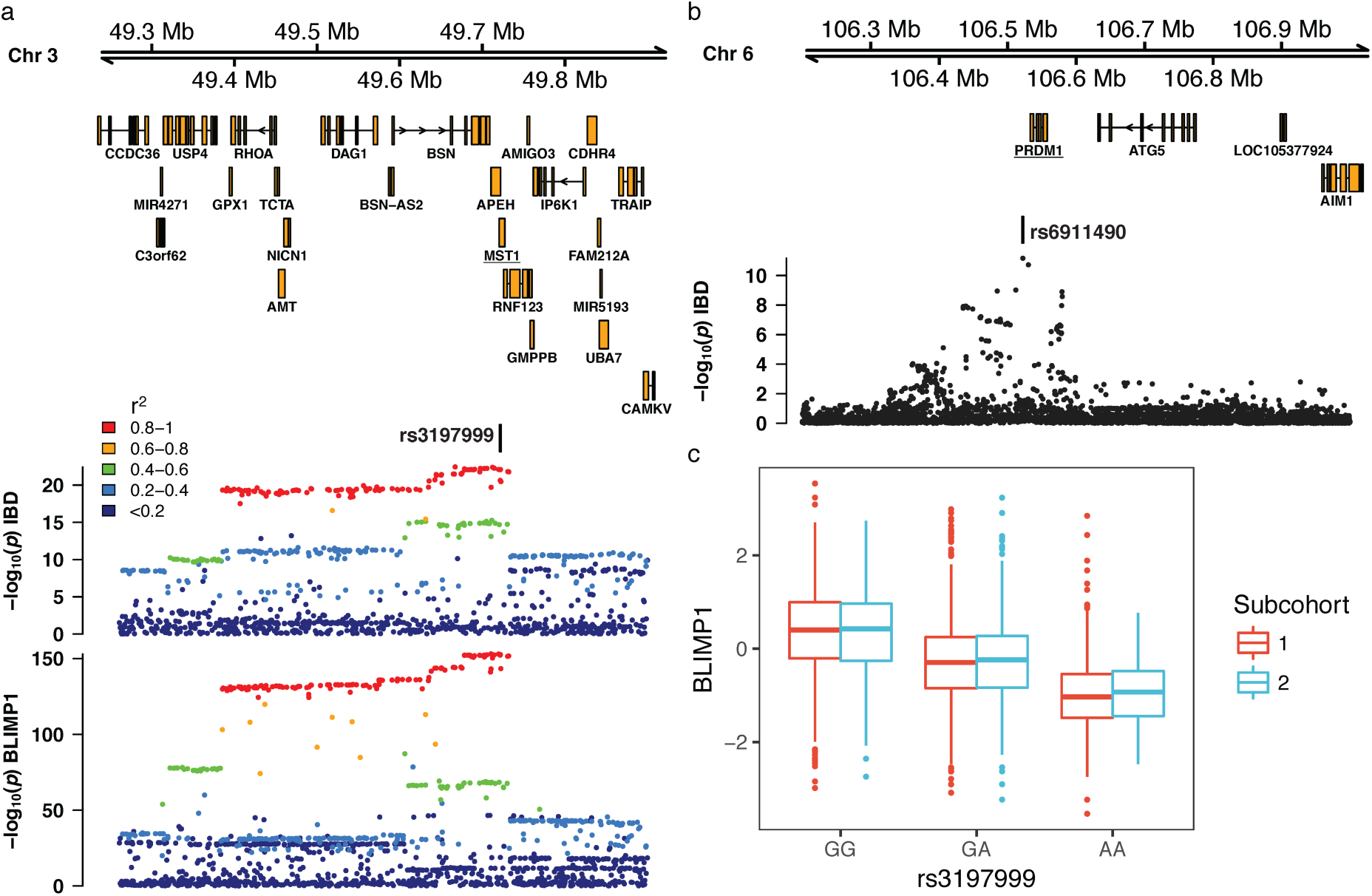
*Trans* pQTL for BLIMP1 at an inflammatory bowel disease (IBD) associated genetic variant in *MST1*. (a) Missense variant rs3197999 in the *MST1* region on chromosome 3 is associated with IBD (top) and BLIMP1 levels (bottom). (b) Regional association plot of the IBD susceptibility locus on chromosome 6 adjacent to the *PRDM1* gene, which encodes BLIMP1. IBD association data are for European participants from Liu *et al.,* 2015. (c) Boxplot for relative plasma BLIMP1 levels by rs3197999 genotype, stratified by the two subcohorts used in our analysis.

We next show how pQTL data can elucidate pathogenic mechanisms. Anti-neutrophil cytoplasmic antibody-associated vasculitis (AAV) is an autoimmune disease characterised by autoantibodies to the neutrophil proteases proteinase-3 (PR3) or myeloperoxidase (MPO). Clinico-pathological features mirror antibody specificity, with granulomatous inflammation typically correlating with anti-PR3 antibodies (PR3+ AAV). GWAS have identified signals in *PRTN3* (encoding PR3) and *SERPINA1* (encoding alpha1-antitrypsin, an inhibitor of PR3) specific to PR3+ AAV^32^. We identified a *cis* pQTL immediately upstream of *PRTN3* (Supplementary Table 1). By linking the risk allele at *PRTN3* to higher plasma levels of the autoantigen PR3, our data strongly suggest a pathogenic role of anti-PR3 antibodies in this disease.

The vasculitis risk allele at the *SERPINA1* locus (rs28929474:T, also known as the ‘Z’ allele) is a missense variant (p.Glu366Lys) which results in defective secretion of alpha1-antitrypsin. We found that the Z allele was not only associated with lower plasma alpha1-antitrypsin, but was also a pQTL “hotspot” associated with 13 proteins using our conservative significance threshold (*p*<1.5×10^−11^) (Figure 4) and 19 at *p*<5×10^−8^. This finding illustrates how a single mutation can lead to widespread perturbation of downstream proteins. One of these proteins was NCF2 (neutrophil cytosolic factor 2), which plays a key role in the neutrophil oxidative burst. Mutations in *NCF2* can result in a rare condition known as chronic granulomatous disease, which like PR3+ AAV exhibits granulomatous inflammation. Our results suggest NCF2 may mediate this characteristic feature of PR3+ AAV.

**Figure 4.**
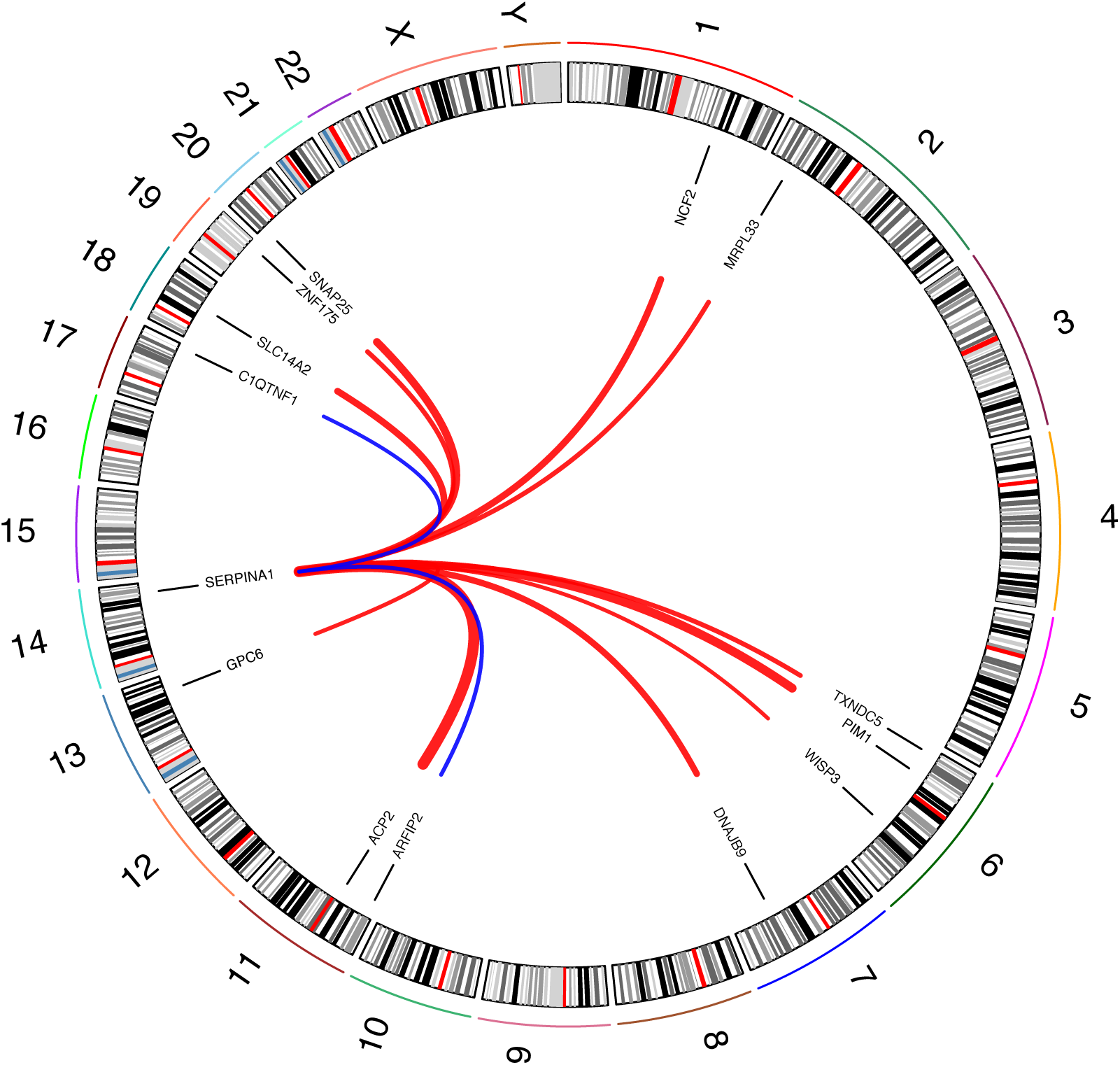
Missense variant rs28929474 in *SERPINA1* is a *trans* pQTL hotspot. Numbers (outermost) indicate chromosomes. Interconnecting lines link the genomic location of rs28929474 and the genes encoding significantly associated (*p*<1.5× 10^−11^) proteins. Line thickness is proportional to the effect size of the associations with red positive and blue negative.

### Causal evaluation of candidate proteins in disease

Association of plasma protein levels with disease does not necessarily imply causation. To help establish causality, we employed the principle of Mendelian randomization (MR)^33^. In contrast with observational studies, which are liable to confounding and/or reverse causation, MR analysis is akin to a “natural” randomised controlled trial, exploiting the random allocation of alleles at conception (Extended Data Figure 6). Consequently, if a genetic variant that specifically influences levels of a protein is also associated with disease risk, then it provides strong evidence of the protein’s causal role. For example, serum levels of PSP-94 (MSMB), a hormone synthesized by the prostate gland and secreted into seminal fluid, are lower in patients with prostate cancer; PSP-94 is therefore a candidate biomarker for prostate cancer^34^. However, it is debated whether PSP-94 plays a causal role in tumorigenesis. We identified a *cis* pQTL for PSP-94 at a prostate cancer susceptibility locus^35^ and showed that the risk allele is associated with lower PSP-94 plasma levels, supporting a protective role for the protein.

In an approach that extends classic MR analysis, we leveraged multi-variant MR analysis methods to distinguish causal genes among multiple plausible candidates at disease loci (Methods), exemplified by the *IL1RL1-IL18R1* locus. We identify four proteins that each had *cis* pQTLs at this locus (Supplementary Table 1), which has been associated with a range of immune-mediated diseases including atopic dermatitis^36^. We created a genetic score for each protein using multiple protein-increasing alleles. Initial “one-protein-at-a-time” analysis identified associations of the scores for IL18R1 (*p=*9.3×10^−72^) and IL1RL1 (*p=*5.7×10^−27^) with atopic dermatitis risk (Figure 5a). In addition, we found a weak association for IL1RL2 (*p=*0.013). We then mutually adjusted these associations for one another to account for the effects of the variants on multiple proteins. While the association of IL18R1 remained significant (*p=*1.5x10^−28^), the association of IL1RL1 (*p=*0.01) was attenuated. In contrast, the association of IL1RL2 (*p*=1.1×10^−69^) became much stronger, suggesting that IL1RL2 and IL18R1 are the causal proteins influencing risk of atopic dermatitis at this locus.

**Figure 5.**
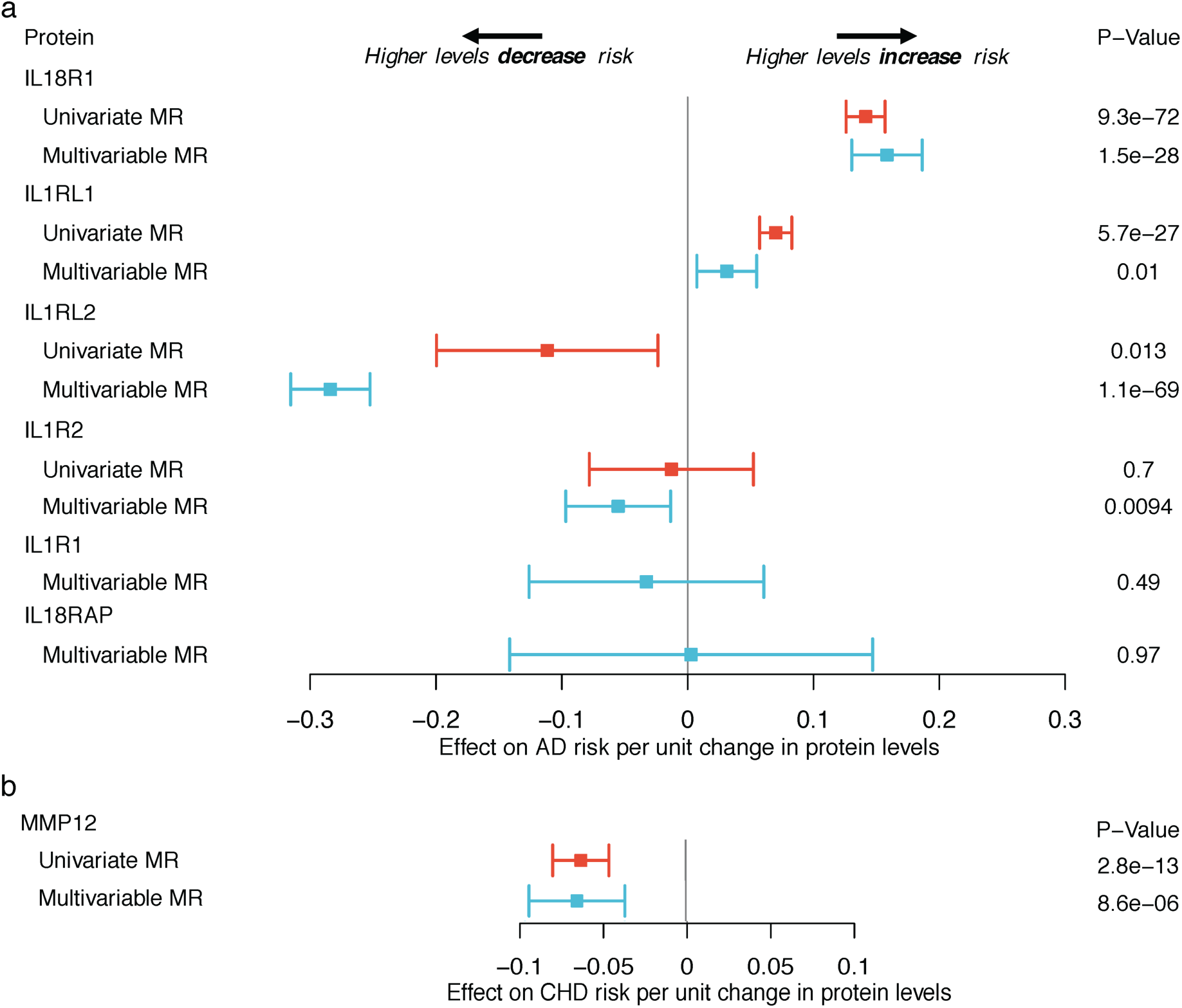
Evaluation of causal role of proteins in disease. Forest plot of univariate and multivariable Mendelian randomization (MR) estimates. (a) Proteins in the *IL1RL1-IL18R1* locus and risk of atopic dermatitis (AD). No univariate MR estimates available for IL1R1 and IL18RAP due to no significant pQTLs to select as a “genetic instrument”. (b) MMP-12 levels and risk of coronary heart disease (CHD).

We also used multiple *cis*-associated variants to evaluate whether macrophage metalloelastase (MMP-12) plays a causal role in coronary disease. Since MMP-12 is known to play a role in lung damage, MMP-12 inhibitors are being tested in chronic obstructive pulmonary disease (COPD). Observational studies report associations of higher levels of plasma MMP-12 with recurrent cardiovascular events^37,38^. In contrast, our multi-allelic genetic score, which explains 14% of the variation in plasma MMP-12 levels (Methods), indicates that genetic predisposition to higher MMP-12 levels is associated with *decreased* coronary disease risk (*p=*2.8×10^−13^) (Figure 5b) and *decreased* large artery atherosclerotic stroke risk^39^. As MMP-12 is released from macrophages in response to cardiovascular injury^40^, it is possible that higher MMP-12 levels are cardioprotective. Hence, our results identify a potential cardiovascular safety concern for MMP-12 inhibitors, particularly as patients with COPD are at high baseline cardiovascular risk due to smoking history.

### Drug target prioritisation

Because therapeutic targets implicated by human genetic data are likely to play causal roles in disease, drugs directed at such targets have greater likelihood of success^41^. According to the Informa Pharmaprojects database (Citeline), 244 of the proteins linked to a disease-associated variant by our pQTL data are drug targets (Supplementary Table 11). Of the proteins we identified as associated with disease susceptibility loci, 49 are targets of already approved drugs such as tocilizumab and ustekinumab (Supplementary Table 12).

To identify additional indications for existing drugs, we investigated disease associations of pQTLs for proteins already targeted by licensed drugs. Our results suggest potential drug “re-purposing” opportunities (Supplementary Table 12). For example, we identified a *cis* pQTL for RANK (encoded by *TNFRSF11A*) at a locus associated with Paget’s disease^42^, a condition characterised by excessive bone turnover leading to deformity and fracture. Standard treatment consists of osteoclast inhibition with bisphosphonates, originally developed as anti-osteoporotic drugs. Denosumab, another anti-osteoporosis drug, is a monoclonal antibody targeting RANKL, the ligand for RANK. Our data suggest denosumab may be a useful alternative for patients with Paget’s disease in whom bisphosphonates are contra-indicated, a hypothesis supported by clinical case reports^43,44^.

To evaluate targets for drugs currently under development, we considered the example of glycoprotein Ib platelet alpha subunit (GP1BA), the receptor for von Willebrand factor. Drugs directed at GP1BA are in pre-clinical development as anti-thrombotic agents and in phase 2 trials to treat thrombotic thrombocytopenic purpura, a life-threatening disorder. We identify a *trans* pQTL for GP1BA at the pleiotropic *SH2B3/BRAP* locus, which is associated with platelet count^45^, myocardial infarction and stroke. The risk allele for cardiovascular disease increases both plasma GP1BA and platelet count, suggesting a mechanism by which this locus affects disease susceptibility. As a confirmation of the link between GP1BA and the platelet count, we found a directionally concordant *cis* pQTL for GP1BA at a platelet count-associated variant (Supplementary Table 12). Collectively, these results suggest that targeting GP1BA may be efficacious in conditions characterised by platelet aggregation such as arterial thrombosis. More generally, our data provide a foundation for generating hypotheses about targets for new drug development through the approach of linking genetic factors to disease via specific proteins (Supplementary Table 12).

## Discussion

We conducted protein measurements of unprecedented scope and scale to reveal genetic control of the human plasma proteome. Our discoveries enabled identification of important consequences of natural perturbations in the plasma proteome. First, we identified the downstream effects of alterations in specific protein levels, revealing novel regulators of protein pathways. Second, we used plasma proteins to uncover intermediate molecular pathways that connect the genome to disease endpoints. Previous investigation has focused on genes in the vicinity of disease susceptibility loci. By contrast, we made key advances through use of *trans* pQTLs to implicate previously unsuspected proteins encoded by distant genes. Third, we established causal roles for protein biomarkers in vascular, neoplastic, and autoimmune diseases using the principle of Mendelian randomization (MR). Proteins provide an ideal paradigm for MR analysis because they are under proximal genetic control. Whereas MR studies of plasma proteins have been constrained by availability of few suitable genetic instruments, our data remedy this bottleneck by furnishing an extensive toolkit. Fourth, we introduced an approach that should help reduce the unsustainably high attrition rates of drugs in pharmaceutical pipelines. Overall, our study foreshadows major advances in post-genomic science through increasing application of novel bioassay technologies to major population biobanks.

## Methods

### Study participants

The INTERVAL study comprises approximately 50,000 participants nested within a randomised trial of blood donation intervals^46^. Between mid-2012 and mid-2014, whole-blood donors aged 18 years and older were consented and recruited at 25 centers of England’s National Health Service Blood and Transplant. All participants completed an online questionnaire including questions about demographic characteristics (e.g., age, sex, ethnic group), anthropometry (height, weight), lifestyle (e.g., alcohol and tobacco consumption) and diet. Participants were generally in good health because blood donation criteria exclude people with a history of major disease (such as myocardial infarction, stroke, cancer, HIV, and hepatitis B or C) and those who have had recent illness or infection. For protein assays, we randomly selected two non-overlapping subcohorts of 2,731 and 831 participants from INTERVAL. Participant characteristics are shown in Supplementary Table 13.

### Plasma sample preparation

Sample collection procedures for INTERVAL have been described previously^46^. In brief, blood samples for research purposes were collected in 6ml EDTA tubes using standard venepuncture protocols. The tubes were inverted three times and transferred at room temperature to UK Biocentre (Stockport, UK) for processing. Plasma was extracted into two 0.8ml plasma aliquots by centrifugation and subsequently stored at -80°C prior to use.

### Protein measurements

We used a multiplexed, aptamer-based approach (SOMAscan assay) to measure the relative concentrations of 3,620 plasma proteins/protein complexes assayed using 4,034 aptamers (“SOMAmer reagents”, hereafter referred to as “SOMAmers”). The assay extends the lower limit of detectable protein abundance afforded by conventional approaches (e.g., immunoassays), measuring both extracellular and intracellular proteins (including soluble domains of membrane-associated proteins), with a bias towards proteins likely to be found in the human secretome (Extended Data Figure 7a)^6,47^. The proteins cover a wide range of molecular functions (Extended Data Figure 7b). The selection of proteins on the platform reflects both the availability of purified protein targets and a focus on proteins known to be involved in pathophysiology of human disease.

Aliquots of 150μl of plasma were sent on dry ice to SomaLogic Inc. (Boulder, Colorado, US) for protein measurement. Assay details have been previously described^47–49^ and a technical white paper with further information can be found at the manufacturer’s website (http://info.somalogic.com/lp_campaign_somascan-white-paper_jan-2016). In brief, modified single-stranded DNA SOMAmers are used to bind to specific protein targets that are then quantified using a DNA microarray. Protein concentrations are quantified as relative fluorescent units.

Quality control (QC) was performed at the sample and SOMAmer level using control aptamers, as well as calibrator samples. At the sample level, hybridization controls on the microarray were used to correct for systematic variability in hybridization, while the median signal over all features assigned to one of three dilution sets (40%, 1% and 0.005%) was used to correct for within-run technical variability. The resulting hybridization scale factors and median scale factors were used to normalise data across samples within a run. The acceptance criteria for these values are between 0.4 and 2.5 based on historical runs. SOMAmer-level QC made use of replicate calibrator samples using the same study matrix (plasma) to correct for between-run variability. The acceptance criteria for each SOMAmer is that the calibration scale factor be less than 0.4 from the median for each of the plates run. In addition, at the plate level, the acceptance criteria are that the median of the calibration scale factors is between 0.8 and 1.2, while 95% of individual SOMAmers must be less than 0.4 from the median within the plate.

In addition to QC processes routinely conducted by SomaLogic, we measured protein levels of 30 and 10 pooled plasma samples randomly distributed across plates for subcohort 1 and subcohort 2, respectively. This approach, which involved masking laboratory technicians to the presence of pooled samples, enabled estimation of the reproducibility of the protein assays in directly relevant samples. We calculated coefficients of variation (CVs) for each SOMAmer within each subcohort by dividing the standard deviation by the mean of the pooled plasma sample protein readouts. In addition to passing SomaLogic QC processes, we required SOMAmers to have a CV ≤ 20% in both subcohorts. Eight non-human protein targets were also excluded, leaving 3,283 SOMAmers (mapping to 2,994 unique proteins/protein complexes) for inclusion in the GWAS.

Protein mapping to UniProt identifiers and gene names was provided by SomaLogic. Mapping to Ensembl gene IDs and genomic positions was performed using Ensembl Variant Effects Predictor v83 (VEP)^50^. Protein subcellular locations were determined by exporting the subcellular location annotations from UniProt^51^. If the term “membrane” was included in the descriptor, the protein was considered to be a membrane protein, whereas if the term “secreted” (but not “membrane”) was included in the descriptor, the protein was considered to be a secreted protein. Proteins not annotated as either membrane or secreted proteins were classified (by inference) as intracellular proteins. Proteins were mapped to molecular functions using gene ontology annotations^52^ from UniProt.

### Non-genetic associations of proteins

To validate the protein assays, we attempted to replicate the associations with age or sex of 45 proteins previously reported by Ngo *et al* and 40 reported by Menni *et al*^48,53^. We used Bonferroni-corrected *p*-value thresholds of *p*=1.1×10^−3^ (0.05/45) and *p*=1.2×10^−3^ (0.05/40) respectively. Relative protein abundances were rank-inverse normalised within each subcohort and linear regression was performed using age, sex, BMI, natural log of estimated glomerular filtration rate (eGFR) and subcohort as independent variables.

### Genotyping and imputation

The genotyping protocol and QC for the INTERVAL samples (n∼50,000) have been described previously in detail^14^. Briefly, DNA extracted from buffy coat was used to assay approximately 830,000 variants on the Affymetrix Axiom UK Biobank genotyping array at Affymetrix (Santa Clara, California, US). Genotyping was performed in multiple batches of approximately 4,800 samples each. Sample QC was performed including exclusions for sex mismatches, low call rates, duplicate samples, extreme heterozygosity and non-European descent. An additional exclusion made for this study was of one participant from each pair of close (first- or second-degree) relatives, defined as 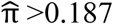. Identity-by-descent was estimated using a subset of variants with call rate >99% and MAF >5% in the merged dataset of both subcohorts, pruned for linkage disequilibrium (LD) using PLINK v1.9^54^. Numbers of participants excluded at each stage of the genetic QC are summarized in Extended Data Figure 1. Multi-dimensional scaling was performed using PLINK v1.9 to create components to account for ancestry in genetic analyses.

Prior to imputation, additional variant filtering steps were performed to establish a high quality imputation scaffold. In summary, 654,966 high quality variants (autosomal, non-monomorphic, bi-allelic variants with Hardy Weinberg Equilibrium (HWE) *p*>5×10^−6^, with a call rate of >99% across the INTERVAL genotyping batches in which a variant passed QC, and a global call rate of >75% across all INTERVAL genotyping batches) were used for imputation. Variants were phased using SHAPEIT3 and imputed using a combined 1000 Genomes Phase 3-UK10K reference panel. Imputation was performed via the Sanger Imputation Server (https://imputation.sanger.ac.uk) resulting in 87,696,888 imputed variants.

Prior to genetic association testing, variants were filtered in each subcohort separately using the following exclusion criteria: (1) imputation quality (INFO) score<0.7, (2) minor allele count<8, (3) HWE *p*<5×10^−6^. In the small number of cases where imputed variants had the same genomic position (GRCh37) and alleles, the variant with the lowest INFO score was removed. 10,572,788 variants passing all filters in both subcohorts were taken forward for analysis (Extended Data Figure 1).

### Genome-wide association study

Within each subcohort, relative protein abundances were first natural log-transformed. Log transformed protein levels were then adjusted in a linear regression for age, sex, duration between blood draw and processing (binary, ≤1 day/>1day) and the first three principal components of ancestry from multi-dimensional scaling. The protein residuals from this linear regression were then rank-inverse normalized and used as phenotypes for association testing. Univariate linear regression using an additive genetic model was used to test genetic associations. Association tests were carried out on allelic dosages to account for imputation uncertainty (“-method expected” option) using SNPTEST v2.5.2^55^.

### Meta-analysis and statistical significance

Association results from the two subcohorts were combined via fixed-effects inverse-variance meta-analysis combining the betas and standard errors using METAL^56^. Genetic associations were considered to be genome-wide significant based on a conservative strategy requiring associations to have (i) a meta-analysis *p*-value<1.5×10^−11^ (genome-wide threshold of *p=*5×10^−8^ Bonferroni corrected for 3,283 aptamers tested), (ii) at least nominal significance (*p<*0.05) in both subcohorts, and also (iii) consistent direction of effect across subcohorts. We did not observe significant genomic inflation (mean inflation factor was 1.0, standard deviation=0.01) (Extended Data Figure 8).

### Refinement of significant regions

To identify distinct non-overlapping regions associated with a given SOMAmer, we first defined a 1Mb region around each significant variant for that SOMAmer. Starting with the region containing the variant with the smallest *p*-value, any overlapping regions were then merged and this process was repeated until no more overlapping 1Mb regions remained. The variant with the lowest *p*-value for each region was assigned as the “regional sentinel variant”. Due to the complexity of the Major Histocompatibility Region (MHC) region, we treated the extended MHC region (chr6:25.5-34.0Mb) as one region. To identify whether a region was associated with multiple SOMAmers, we used an LD-based clumping approach. Regional sentinel variants in high LD (*r*^2^≥0.8) with each other were combined together into a single region.

### Conditional analyses

To identify conditionally significant signals, we performed approximate genome-wide step-wise conditional analysis using GCTA v1.25.2^57^ using the “cojo-slct” option. We used the same conservative significance threshold of *p*=1.5×10^−11^ as for the univariate analysis. As inputs for GCTA, we used the summary statistics (i.e. betas and standard errors) from the meta-analysis. Correlation between variants was estimated using the hard-called genotypes (where a genotype was called if it had a posterior probability of >0.9 following imputation or set to missing otherwise) in the merged genetic dataset, and only variants also passing the univariate genome-wide threshold (*p<*1.5×10^−11^) were considered for step-wise selection. As the conditional analyses use different data inputs (summarised rather than individual-level data), there were some cases where the conditional analysis failed to include sentinel variants that had borderline significant univariate associations in the step-wise selection. In these instances (n=28), we re-conducted the joint model estimation without step-wise selection in GCTA with variants identified by the conditional analysis in addition to the regional sentinel variant. We report and highlight these cases in Supplementary Table 2.

### Replication of previous pQTLs

We attempted to identify all previously reported pQTLs from GWAS and to assess whether they replicated in our study. We used the NCBI Entrez programming utility in R (rentrez) to perform a literature search for pQTL studies published from 2008 onwards. We searched for the following terms: “pQTL”, “pQTLs”, and “protein quantitative trait locus”. We supplemented this search by filtering out GWAS associations from the NHGRI-EBI GWAS Catalog v.1.0.1^58^ (https://www.ebi.ac.uk/gwas/, downloaded April 2016), which has all phenotypes mapped to the Experimental Factor Ontology (EFO)^59^, by restricting to those with EFO annotations relevant to protein biomarkers (e.g. “protein measurement”, EFO_0004747). Studies identified through both approaches were manually filtered to include only studies that profiled plasma or serum samples and to exclude studies not assessing proteins. We recorded basic summary information for each study including the assay used, sample size and number of proteins with pQTLs (Supplementary Table 14). To reduce the impact of ethnic differences in allele frequencies on replication rate estimates, we filtered studies to include only associations reported in European-ancestry populations. We then manually extracted summary data on all reported associations from the manuscript or the supplementary material. This included rsID, protein UniProt ID, *p*-values, and whether the association is *cis/trans* (Supplementary Table 15).

To assess replication we first identified the set of unique UniProt IDs that were also assayed on the SOMAscan panel. For previous studies that used SomaLogic technology, we refined this match to the specific aptamer used. We then clumped associations into distinct loci using the same method applied to our pQTL (see **Refinement of significant regions**). For each locus, we asked if the sentinel SNP or a proxy (*r*^2^>0.6) was associated with the same protein/aptamer in our study at a defined significance threshold. For our primary assessment, we used a *p*-value threshold of 10^−4^. We also performed sensitivity analyses to explore factors that influence replication rate (Supplementary Note).

### Candidate gene annotation

We defined a pQTL as *cis* when the most significantly associated variant in the region was located within 1Mb of the transcription start site (TSS) of the gene(s) encoding the protein. pQTLs lying outside of the region were defined as *trans*. When considering the distance of the top *cis*-associated variant from the relevant TSS, only proteins that map to single genes on the primary assembly in Ensembl v83 were considered.

For *trans* pQTLs, we sought to prioritise candidate genes in the region that might underpin the genotype-protein association. In addition to reporting the nearest gene to the sentinel variant, we employed “bottom up” and “top down” approaches, starting from the variant and protein respectively For the “bottom up” approach, the sentinel variant and corresponding proxies (*r*^2^>0.8) for each *trans* pQTL were first annotated using Ensembl VEP v83 (using the “pick” option) to determine whether variants were (1) protein-altering coding variants; (2) synonymous coding or 5’/3’ untranslated region (UTR); (3) intronic or up/downstream; or (4) intergenic. Second, we queried all sentinel variants and proxies against significant *cis* eQTL variants (defined by beta distribution-adjusted empirical *p*-values using a FDR threshold of 0.05, see http://www.gtexportal.org/home/documentationPage for details) in any cell type or tissue from the Genotype-Tissue Expression (GTEx) project v6^60^ (http://www.gtexportal.org/home/datasets). Third, we also queried promoter capture Hi-C data in 17 human primary hematopoietic cell types^61^ to identify contacts (with a CHICAGO score >5 in at least one cell type) involving chromosomal regions containing a sentinel variant. We considered gene promoters annotated on either fragment (i.e., the fragment containing the sentinel variant or the other corresponding fragment) as potential candidate genes. Using these three sources of information, we generated a list of candidate genes for the *trans* pQTLs. A gene was considered a candidate if it fulfilled at least one of the following criteria: (1) it was proximal (intragenic or ±5Kb from the gene) or nearest to the sentinel variant; (2) it contained a sentinel or proxy variant (*r*^2^>0.8) that was protein-altering; (3) it had a significant *cis* eQTL in at least one GTEx tissue overlapping with a sentinel pQTL variant (or proxy); or (4) it was regulated by a promoter annotated on either fragment of a chromosomal contact^61^ involving a sentinel variant.

For the “top down” approach, we first identified all genes with a TSS located within the corresponding pQTL region using the GenomicRanges Bioconductor package^62^ with annotation from a GRCh37 GTF file from Ensembl (ftp://ftp.ensembl.org/pub/grch37/update/gtf/homo_sapiens/; file: “Homo_sapiens.GRCh37.82.gtf.gz”, downloaded June 2016). We then identified any local genes that had previously been linked with the corresponding *trans*-associated protein(s) according to the following open source databases: (1) the Online Mendelian Inheritance in Man (OMIM) catalogue^63^ (http://www.omim.org/); (2) the Kyoto Encyclopedia of Genes and Genomes (KEGG)^64^ (http://www.genome.jp/kegg/); and (3) STRINGdb^65^ (http://string-db.org/; v10.0). We accessed OMIM data via HumanMine web tool^66^ (http://www.humanmine.org/; accessed June 2016), whereby we extracted all OMIM IDs for (i) our *trans*-affected proteins and (ii) genes local (±500Kb) to the corresponding *trans*-acting variant. We extracted all human KEGG pathway IDs using the KEGGREST Bioconductor package^67^ (https://bioconductor.org/packages/release/bioc/html/KEGGREST.html). In cases where a *trans*-associated protein shared either an OMIM ID or a KEGG pathway ID with a gene local to the corresponding *trans*-acting variant, we took this as evidence of a potential functional involvement of that gene. We interrogated protein-protein interaction data by accessing STRINGdb data using the STRINGdb Bioconductor package^68^, whereby we extracted all pairwise interaction scores for each *trans*-affected protein and all proteins with genes local to the corresponding *trans*-acting variants. We took the default interaction score of 400 as evidence of an interaction between the proteins, therefore indicating a possible functional involvement for the local gene. In addition to using data from open source databases in our top-down approach we also adopted a “guilt-by-association” (GbA) approach utilising the same plasma proteomic data used to identify our pQTLs. We first generated a matrix containing all possible pairwise Pearson’s correlation coefficients between our 3,283 SOMAmers. We then extracted the coefficients relating to our *trans-*associated proteins and any proteins encoded by genes local to their corresponding *trans*-acting variants (where available). Where the correlation coefficient was ≥0.5 we prioritised the relevant local genes as being potential mediators of the *trans*-signal(s) at that locus.

We report the potential candidate genes for our *trans* pQTLs from both the “bottom up” and “top down” approaches, highlighting cases where the same gene was highlighted by both approaches.

### Functional annotation of pQTLs

Functional annotation of variants was performed using Ensembl VEP v83 using the “pick” option. We tested the enrichment of significant pQTL variants for certain functional classes by comparing to permuted sets of variants showing no significant association with any protein (*p*>0.0001 for all proteins tested). First the regional sentinel variants were LD-pruned at *r*^2^ of 0.1. Each time the sentinel variants were LD-pruned, one of the pair of correlated variants was removed at random and for each set of LD-pruned sentinel variants and 100 sets of equally sized null permuted variants were sampled matching for MAF (bins of 5%), distance to TSS (bins of 0-0.5Kb, 0.5Kb-2Kb, 2Kb-5Kb, 10Kb-20Kb, 20Kb-100Kb and >100Kb in each direction) and LD (± half the number of variants in LD with the sentinel variant at *r*^2^ of 0.8). This procedure was repeated 100 times resulting in 10,000 permuted sets of variants. An empirical *p*-value was calculated as the proportion of permuted variant sets where the proportion that are classified as a particular functional group exceeded that of the test set of sentinel pQTL variants, and we used a significance threshold of *p*=0.005 (0.05/10 functional classes tested).

### Evidence against aptamer-binding effects at *cis* pQTLs

All protein assays that rely on binding (e.g. of antibodies or SOMAmers) are susceptible to the possibility of binding-affinity effects, where protein-altering variants (PAVs) (or their proxies in LD) are associated with protein measurements due to differential binding rather than differences in protein abundance. To account for this potential effect, we performed conditional analysis at all *cis* pQTLs where the sentinel variant was in LD (0.1≤*r*^2^≤0.9) with a PAV in the gene(s) encoding the associated protein. First, variants were annotated with Ensembl VEP v83 using the “per-gene” option. Variant annotations were considered protein altering if they were annotated as coding sequence variant, frameshift variant, in-frame deletion, in-frame insertion, missense variant, protein altering variant, splice acceptor variant, splice donor variant, splice region variant, start lost, stop gained, or stop lost. To avoid multi-collinearity, PAVs were LD-pruned (*r*^2^>0.9) using PLINK v1.9 before including them as covariates in the conditional analysis on the meta-analysis summary statistics using GCTA v1.25.2. Any variants with an *r*^2^≥0.9 with any of the PAVs were removed. Coverage of known common (MAF>5%) PAVs in our data was checked by comparison with exome sequences from ∼60,000 individuals in the Exome Aggregation Consortium^69^ (ExAC [http://exac.broadinstitute.org], downloaded June 2016).

### Testing for regulatory and functional enrichment

We tested whether our pQTLs were enriched for functional and regulatory characteristics using GARFIELD v1.2.0^18^. GARFIELD is a non-parametric permutation-based enrichment method that compares input variants to permuted sets matched for number of proxies (*r*^2^≥0.8), MAF and distance to the closest TSS. It first applies “greedy pruning” (*r*^2^<0.1) within a 1Mb region of the most significant variant. GARFIELD annotates variants with more than a thousand features, drawn predominantly from the GENCODE, ENCODE and ROADMAP projects, which includes genic annotations, histone modifications, chromatin states and other regulatory features across a wide range of tissues and cell types.

The enrichment analysis was run using all variants that passed our Bonferroni-adjusted significance threshold *(p<*1.5×10^−11^) for association with any protein. For each of the matching criteria (MAF, distance to TSS, number of LD proxies), we used five bins. In total we tested 25 combinations of features (classified as transcription factor binding sites, FAIRE-seq, chromatin states, histone modifications, footprints, hotspots, or peaks) with up to 190 cell types from 57 tissues, leading to 998 tests. Hence, we considered enrichment with a *p*<5 ×10^−5^ (0.05/998) to be statistically significant.

### Disease annotation

To identify diseases that our pQTLs have been associated with, we queried our sentinel variants and their strong proxies (*r*^2^≥0.8) against publicly available disease GWAS data using PhenoScanner^8^. A list of datasets queried is available at http://www.phenoscanner.medschl.cam.ac.uk/information.html. For disease GWAS, results were filtered to *p*<5x10^−8^ and then manually curated to retain only the entry with the strongest evidence for association (i.e. smallest *p*-value) per disease. Non-disease phenotypes such as anthropometric traits, intermediate biomarkers and lipids were excluded manually.

### *Cis* eQTL overlap and enrichment of *cis* pQTLs for *cis* eQTLs

Each regional sentinel *cis* variant its strong proxies (*r*^2^≥0.8) were queried against publicly available eQTL association data using PhenoScanner. *Cis* eQTL results were filtered to retain only variants with *p*<1.5×10^−11^. Only *cis* eQTLs for the same gene as the *cis* pQTL protein were retained. To assess whether our *cis* pQTLs were more likely to also be *cis* eQTLs than non-pQTL variants, we used data from the GTEx project v6, due to the availability of genome-wide association results across a wide range of tissues and cell-types. GTEx results were filtered to contain only variants lying in *cis* (i.e., within 1Mb) of genes that encode proteins analysed in our study and only variants in both datasets were utilised.

For the enrichment analysis, the *cis* pQTL sentinel variants were first LD-pruned (*r*^2^ <0.1) and the proportion of sentinel *cis* pQTL variants that are also eQTLs (at *p*<1.5×10^−11^) for the same protein/gene was compared to a permuted set of variants that were not pQTLs *(p*>0.0001 for all proteins). We generated 10,000 permuted sets of null variants matched for MAF, distance to TSS and LD (as described for functional annotation enrichment in **Functional annotation of pQTLs**). An empirical *p*-value was calculated as the proportion of permuted variant sets where the proportion that are also *cis* eQTLs exceeded that of the test set of sentinel *cis* pQTL variants. Results were similar in sensitivity analyses using the standard genome-wide significance threshold of *p*<5×10^−8^ for the eQTLs (13.0% for whole blood, 18.7% for LCLs, 17.1% for liver and 13.4% for monocytes) as well as also using only the sentinel variants at *cis* pQTLs that were robust to adjusting for PAVs, suggesting our results are robust to choice of threshold and potential differential binding effects.

### Mediation of the GDF11/8 *trans* pQTL by WFIKKN2 levels

To assess whether the *trans* pQTL for GDF11/8 in the *WFIKKN2* gene region was mediated by WFIKKN2 levels, we first regressed the rank inverse transformed residuals for GDF11/8 used for the GWAS against the WFIKKN2 residuals, adjusting for subcohort. The residuals from this regression were subsequently regressed against allelic dosages for each variant in the *WFIKKN2* region. As there were two SOMAmers targeting WFIKKN2, we tested both to see if similar results were obtained. Regional association plots were made using Gviz^70^.

### Selection of genetic instruments for Mendelian randomization

In Mendelian randomization (MR), genetic variants are used as “instrumental variables” (IV) for assessing the causal effect of the exposure (here a plasma protein) on the outcome (here disease).

### Proteins in the *IL1RL1-IL18R1* locus and atopic dermatitis

To identify the likely causal proteins that underpin the previous genetic association of the *IL1RL1-IL18R1* locus (chr11:102.5-103.5Mb) with atopic dermatitis (AD)^36^, we used a multivariable MR approach. For each protein encoded by a gene in the *IL1RL1-IL18R1* locus, we took genetic variants that had a *cis* association at *p*<1×10^−4^ and ‘LD-pruned’ them at *r*^2^<0.1. We then used these variants as instrumental variables for their respective proteins in univariate MR. For multivariable MR, association estimates for all proteins in the locus were extracted for all instruments. We used PhenoScanner to obtain association statistics for the selected variants in the European-ancestry population of a recent large-scale GWAS meta-analysis^36^. Where the relevant variant was not available, the strongest proxy with *r*^2^≥0.8 was used.

### MMP-12 and coronary heart disease (CHD)

To test whether plasma MMP-12 levels have a causal effect on risk of CHD, we selected genetic variants in the *MMP12* gene region to use as instrumental variables. We constructed a genetic score comprising 17 variants that had a *cis* association with MMP-12 levels at *p*<5x10^−8^ and that were not highly correlated with one another (*r*^2^<0.2). To perform multivariable MR, we used association estimates for these variants with other MMP proteins in the locus (MMP-1, MMP-7, MMP-8, MMP-10, MMP-13). Summary associations for variants in the score with CHD were obtained through PhenoScanner from a recent large-scale 1000 Genomes-based GWAS meta-analysis which consists mostly (77%) of individuals of European ancestry^71^.

### MR analysis

Two-sample univariate MR was performed for each protein separately using summary statistics in the inverse-variance weighted method adapted to account for correlated variants^72,73^. For each of *G* genetic variants (*g* = 1, …, *G*) having per-allele estimate of the association with the protein *β*_*Xg*_ and standard error *σ*_*Xg*_, and per-allele estimate of the association with the outcome (here, AD or CHD) *β*_*Yg*_ and standard error *σ*_*Yg*_, the IV estimate 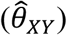 is obtained from generalised weighted linear regression of the genetic associations with the outcome (*β*_*Y*_) on the genetic associations with the protein (*β*_*X*_) weighting for the precisions of the genetic associations with the outcome and accounting for correlations between the variants according to the regression model:

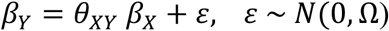

where *β*_*Y*_ and *β*_*X*_ are vectors of the univariable (marginal) genetic associations, and the weighting matrix Ω has terms 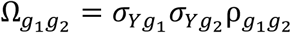, and 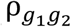 is the correlation between the *g*_1_ th and *g*_2_ th variants.

The IV estimate from this method is:

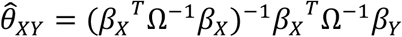

and the standard error is:

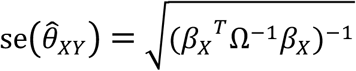

where ^*T*^ is a matrix transpose. This is the estimate and standard error from the regression model fixing the residual standard error to 1 (equivalent to a fixed-effects model in a meta-analysis).

Genetic variants in univariate MR need to satisfy three key assumptions to be valid instruments:

1. the variant is associated with the risk factor of interest (i.e., the protein level),
2. the variant is not associated with any confounder of the risk factor-outcome association,
3. the variant is conditionally independent of the outcome given the risk factor and confounders.

To account for potential effects of functional pleiotropy^74^, we performed multivariable MR using the weighted regression-based method proposed by Burgess *et al*^75^. For each of *K* risk factors in the model (*k* = 1, …, *K*), the weighted regression-based method is performed by multivariable generalized weighted linear regression of the association estimates *β*_*Y*_ on each of the association estimates with each risk factor *β*_*Xk*_ in a single regression model:

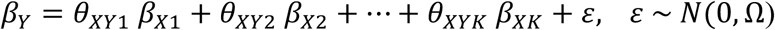

where *β*_*X*1_ is the vectors of the univariable genetic associations with risk factor 1, and so on. This regression model is implemented by first pre-multiplying the association vectors by the Cholesky decomposition of the weighting matrix, and then applying standard linear regression to the transformed vectors. Estimates and standard errors are obtained fixing the residual standard error to be 1 as above.

The multivariable MR analysis allows the estimation of the causal effect of a protein on disease outcome accounting for the fact that genetic variants may be associated with multiple proteins in the region. Causal estimates from multivariable MR represent direct causal effects, representing the effect of intervening on one risk factor in the model while keeping others constant.

### MMP-12 genetic score sensitivity analyses

We performed two sensitivity analyses to determine the robustness of the Mendelian randomization findings. First, we measured plasma MMP-12 levels using a different method (proximity extension assay; Olink Bioscience, Uppsala, Sweden^76,77^) in a sub-sample of 141 individuals, and used this to derive genotype-MMP12 effect estimates for the 17 variants in our genetic score. Second, we obtained effect estimates from a pQTL study based on SOMAscan assay measurements in an independent sample of ∼1,000 individuals^7^. In both cases the genetic score reflecting higher plasma MMP-12 was associated with lower risk of CHD.

### Overlap of pQTLs with drug targets

We used the Informa Pharmaprojects database from Citeline to obtain information on drugs that target proteins assayed on the SOMAscan platform. This is a manually curated database that maintains profiles for >60,000 drugs. For our analysis, we focused on the following information for each drug: protein target, indications, and development status. We included drugs across the development pipeline, including those in pre-clinical studies or with no development reported, drugs in clinical trials (all phases), and launched/registered drugs. For each protein assayed, we identified all drugs in the Informa Pharmaprojects with a matching protein target based on UniProt ID. When multiple drugs targeted the same protein, we selected the drug with the latest stage of development.

For drug targets with significant pQTLs, we identified the subset where the sentinel variant or proxy variants in LD (*r*^2^>0.6) are also associated with disease risk through PhenoScanner. We used an internal Merck auto-encoding method to map GWAS traits and drug indications to a common set of terms from the Medical Dictionary for Regulatory Activities (MedDRA). MedDRA terms are organized into a hierarchy with five levels. We mapped each GWAS trait and indication onto the ‘Lowest Level Terms’ (i.e. the most specific terms available). All matching terms were recorded for each trait or indication. We matched GWAS traits to drug indications based on the highest level of the hierarchy, called ‘System Organ Class’ (SOC). We designated a protein as ‘matching’ if at least one GWAS trait term matched with at least one indication term for at least one drug.

### Data availability

Summary association results will be made available on publication through http://www.phenoscanner.medschl.cam.ac.uk.

## Supplementary Information

Supplementary Information is available in the online version of the paper.

## Acknowledgements

We acknowledge the participation of all INTERVAL volunteers. We thank the INTERVAL study co-ordination teams (at the Universities of Cambridge and Oxford and at NHS Blood and Transplant [NHSBT]), including the blood donation staff at the 25 static centers, for their help with INTERVAL participant recruitment and study fieldwork, as well as the Cambridge BioResource and NHSBT staff for their help with volunteer recruitment. We thank the INTERVAL Operations Team headed by Dr Richard Houghton and Dr Carmel Moore, and the INTERVAL Data Management Team headed by Dr Matthew Walker. We thank all the staff at SomaLogic for processing and running the proteomic assays. We thank Aaron Day-Williams, Joshua McElwee, Dorothee Diogo, William Astle, Emanuele Di Angelantonio, Ewan Birney, Arianne Richard, Justin Mason and Michael Inouye for helpful comments on the manuscript and Mark Sharp for help mapping drug indications to GWAS traits. The MRC/BHF Cardiovascular Epidemiology Unit is supported by the UK Medical Research Council (G0800270), British Heart Foundation (SP/09/002), UK National Institute for Health Research Cambridge Biomedical Research Centre, European Research Council (268834), and European Commission Framework Programme 7 (HEALTH-F2-2012-279233). B.B.S. is funded by the Cambridge School of Clinical Medicine MRC/Sackler Prize PhD Studentship (MR/K50127X/1) and supported by the Cambridge School of Clinical Medicine MB-PhD programme. J.E.P. is funded by a British Heart Foundation Clinical Research Fellowship through the BHF Cambridge Centre of Excellence [RE/13/6/30180]. D.S.P. and D.S. are funded by the Wellcome Trust (105602/Z/14/Z). N.S. is supported by the Wellcome Trust (WT098051 and WT091310), the EU FP7 (EPIGENESYS 257082 and BLUEPRINT HEALTH-F5-2011-282510). J.A.T is supported by the supported by the Wellcome Trust (091157) and JDRF (9-2011-253). K.S. is funded by the Biomedical Research Program funds at Weill Cornell Medicine in Qatar, a program funded by the Qatar Foundation. J.D. is a British Heart Foundation Professor, European Research Council Senior Investigator, and National Institute for Health Research (NIHR) Senior Investigator. The INTERVAL study is funded by NHSBT (11-01-GEN) and has been supported by the NIHR-BTRU in Donor Health and Genomics (NIHR BTRU-2014-10024) at the University of Cambridge in partnership with NHSBT. The views expressed are those of the authors and not necessarily those of the NHS, the NIHR, the Department of Health of England, or NHSBT.

### Author Contributions

Conceptualization and experimental design: J.D., A.S.B., B.B.S., H.R., R.M.P.; Methodology: B.B.S., A.B.S., J.C.M., J.E.P., H.R., S.B.; Analysis: B.B.S., J.C.M., J.E.P., D.S., J.B., J.R.S., T.J., E.P., P.S., C.O-W., M.A.K., S.K.W., A.C., N.B., S.L.S.; Contributed reagents, materials, protocols or analysis tools: N.J., S.K.W., E.S.Z., J.B., M.A.K., J.R.S., B.P.P.; Supervision: A.S.B., H.R., C.S.F., J.D., R.M.P., D.S.P., A.M.W.; Writing - principal: B.B.S., A.S.B., J.E.P., J.C.M., H.R., J.D.; Writing – review and editing: B.B.S., A.S.B., J.E.P., J.C.M., J.D., H.R., K.S., A.M.W., N.J., D.J.R., J.A.T., D.S.P., N.S., C.S.F., R.M.P; Creation of the INTERVAL BioResource: J.R.B., D.J.R., W.H.O., N.W.M., J.D.; Funding acquisition: N.W.M., J.R.B., D.J.R., W.H.O., C.S.F., R.M.P., J.D.; all authors critically reviewed the manuscript.

### Author Information

Reprints and permissions information is available at www.nature.com/reprints. The authors declare the following competing financial interests: J.C.M., A.C., C.S.F., R.M.P., H.R. are employees at MRL, Merck & Co., Inc. S.K.W., E.S.Z., N.J. are employees and stakeholders in SomaLogic, Inc. The other authors have nothing to disclose. Correspondence and requests for materials should be addressed to A.S.B. and J.D.

